# Expression of a mammalian RNA demethylase increases flower number and floral stem branching in *Arabidopsis thaliana*

**DOI:** 10.1101/2024.02.15.580558

**Authors:** Kasey Markel, Lucas Waldburger, Patrick M. Shih

**Affiliations:** Department of Plant and Microbial Biology, University of California, Berkeley, Berkeley, CA, USA; Feedstocks Division, Joint BioEnergy Institute, Emeryville, CA, USA; Environmental Genomics and Systems Biology Division, Lawrence Berkeley National Laboratory, Berkeley, CA, USA; Department of Bioengineering, University of California, Berkeley, California, USA Joint BioEnergy Institute, Emeryville, California, USA; Biological Systems and Engineering Division, Lawrence Berkeley National Laboratory, Berkeley, California, USA; Joint Genome Institute, Lawrence Berkeley National Laboratory, Berkeley, CA, United States; Innovative Genomics Institute, University of California, Berkeley, CA

## Abstract

RNA methylation plays a central regulatory role in plant biology and is a relatively new target for plant improvement efforts. In nearly all cases, perturbation of the RNA methylation machinery results in deleterious phenotypes. However, a recent landmark paper reported that transcriptome-wide use of the human RNA demethylase FTO substantially increased the yield of rice and potatoes. Here, we have performed the first independent replication of those results and broader transferability of the trait, demonstrating increased flower and fruit count in the model species *Arabidopsis thaliana*. We also performed RNA-seq of our FTO-transgenic plants, which we analyzed in conjunction with previously-published datasets to detect several previously-recognized patterns in the functional and structural classification of the upregulated and downregulated genes. From these, we present mechanistic hypotheses to explain these surprising results with the goal of spurring more widespread interest in this promising new approach to plant engineering.

## Introduction

RNA methylation plays a central regulatory role in all eukaryotes, and influences RNA processing, translation rate, and transportation^1^. Over 200 types of RNA modification have been identified in plants, of which the most common is N6-methyladenosine (m^6^A)^2^, which is found on approximately 0.5% of adenosines in mRNA^3^. Most m^6^A in eukaryotes is found within the consensus motif RRACH (R = A/G, H = A/U/C). Enzymes that add the methyl group to adenosine are often called m^6^A writers, enzymes that remove the methyl group are called m^6^A erasers, and proteins that recognize the methyl group are called M^6^A readers. Many plant traits are known to be affected by m^6^A status, including resistance to fungal pathogens^4^ and fruit ripening^5^. Targeted m^6^A editing has been proposed as a method to improve crop quality, with specific genes proposed to target flowering time and fruit maturation^6^.

Recently, Yu *et al*. reported that transgenic expression of the mammalian m^6^A eraser FTO substantially increased yield in both potatoes (*Solanum tuberosum*) and rice (*Oryza sativa*)^7^. They reported an approximately 50% increase in yield in the field, a remarkably large effect size for a single transgene. In addition to the increase in yield, Yu *et al*. reported an increase in shoot biomass and polyadenylated mRNA accumulation. The yield increase was explained through an increase in tillering, which suggests reduced apical dominance and increased branching.

FTO overexpression has been shown to increase biomass across a wide range of animal species. FTO has been intensely studied in humans because it is the QTL most strongly associated with obesity^8,9^. Overexpression of FTO in mice leads to increased food intake and obesity^10^, whereas *fto* mice display reduced growth and lower body mass^8,11^. Overexpression of FTO also increases the mitosis rates of mouse fat cells *in vitro*^9^, and *fto* mutant mice gain less weight even when forced to consume the same amount of food and water as wild-type mice^12^. In porcine cell lines, FTO overexpression results in higher accumulation of lipids while *fto* knockouts accumulate lower levels^13^, and in chicken cells FTO overexpression increases adipogenesis while FTO silencing reduces adipogenesis^14^.

In contrast, there were no reports prior to Yu *et al*. that reduction in m^6^A levels results in increased biomass in plants, though many mutants with altered m^6^A levels have been characterized in the last two decades. In *Arabidopsis*, known m^6^A writers include MTA, MTB, FIP37, VIR, FIONA1, and HAKAI. Complete knockout of MTA is embryo-lethal^15^, but plants rescued with embryo-specific expression display reduced apical dominance and an increase in trichome branch number^16^. Silencing of MTA with artificial microRNA results in overproliferation of shoot apical meristems (SAMs)^17^. Another study found artificial microRNA interference against MTA or VIR resulted in expansion of the SAM onto the petioles of young leaves^18^. RNA interference against MTB also resulted in a reduction in apical dominance and partial dwarfism^19^. *Fip37* knockout lines have reduced apical dominance and severe dwarfism^19^, and rescue with embryo-specific expression resulted in SAM overproliferation^17^. *Fiona1* mutants have altered RNA splicing patterns^20^, early flowering, and more branching in the floral stem^21^. Knockout of the m^6^A eraser ALKBH10B resulted in increased m^6^A, late flowering, and reduced growth, whereas constitutive overexpression resulted in reduced m^6^A and early flowering^22^. While the details vary depending on which gene is perturbed, it is generally the case that reduction of genome-wide m^6^A levels in plants results in reduced apical dominance, earlier flowering, and enlarged meristems. Taken together, there are several lines of evidence suggesting how controlling RNA methylation patterns may provide a means to tune agriculturally relevant traits.

Increasing yield potential through genetic improvement is a major goal of plant biology^23^, and despite massive progress to date, more remains to be accomplished^24^. Given the dramatic nature of the yield increase reported by Yu *et al*., understanding the transferability of this approach into other plants has been of keen interest. Moreover, understanding how RNA methylation plays a role in this change will help elucidate new avenues in crop improvement. Here, we investigated transgenic expression of FTO in *Arabidopsis* and performed RNA-seq to generate hypotheses as to the underlying mechanism.

## Results

### Overexpression of FTO increases many yield-associated traits in *Arabidopsis*

We synthesized an *Arabidopsis* codon-optimized version of the human FTO gene under the control of the pCH3 promoter (At4G13930), which provides high constitutive expression^25^, in a binary plasmid as shown in Figure 1A. We generated *Arabidopsis* stable lines using floral dip and selecting for kanamycin resistance, then grew the plants for two generations and selected for lines with approximately 100% survival on kanamycin, indicating either multiple insertion or homozygosity for FTO. We grew the T3-generation FTO plants and found that they bolted earlier than Col-0 (Figure 1B). Plants were grown an additional 4 days after imaging to allow for maturation and for nearly all plants to bolt, then the plants were destructively phenotyped. Only plants that had bolted (defined as having a floral stem weighing over 100 mg and at least one flower) were included, for a total of 188 plants. Phenotyping consisted of counting the flowers and fruits (siliques), counting the number of shoot tips, and weighing the rosette and floral stems; raw data are available in Supplemental Table 1.

**Figure 1:**
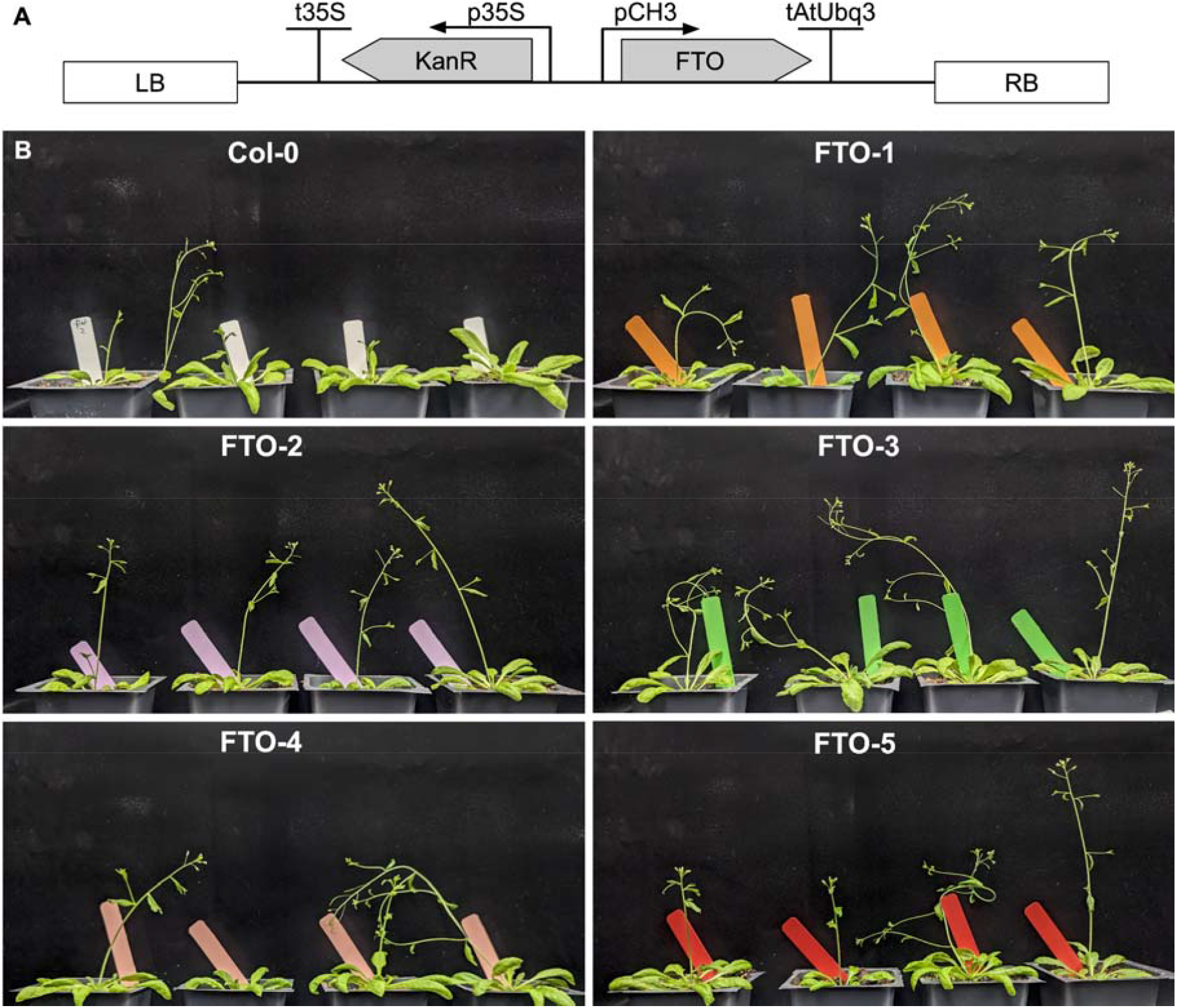
Transgenic expression of FTO causes early bolting in *Arabidopsis*. A) Genetic design of the synthetic FTO construct. B) Transgenic plants of the T3 generation. All plants were grown together and randomly-selected for imaging on the same day from one of the ten flats used in this experiment with no cherry picking.

Though Yu *et al*. claimed a significant increase in stem biomass for both potatoes and rice, we found no difference in *Arabidopsis* rosette mass between FTO plants and Col-0 wild-type (Figure 2A). However, the floral stem mass was significantly increased for all five independent FTO lines (Figure 2B). There was no change in total shoot mass (Figure 2C), suggesting the key difference between FTO and Col-0 plants was the allocation of growth towards sexual rather than vegetative tissue. Accordingly, there was a significant increase in percentage of stem mass apportioned in floral stem tissue (Figure 2D) for four out of five lines. All FTO lines had a larger number of flowers and siliques than Col-0, with an average increase of 100% (Figure 2E). Though *a priori* surprising, this result aligns well with Yu *et al*.’s finding of a ∼200% increase in yield in rice in the greenhouse and ∼50% in the field. Our result in a different species in growth chambers is within the range of their observed increases in yield. The effect was statistically significant for all lines, with a maximum p-value of 0.015 (Wilcoxon test with Benjamini-Hochberg correction for multiple comparisons).

**Figure 2:**
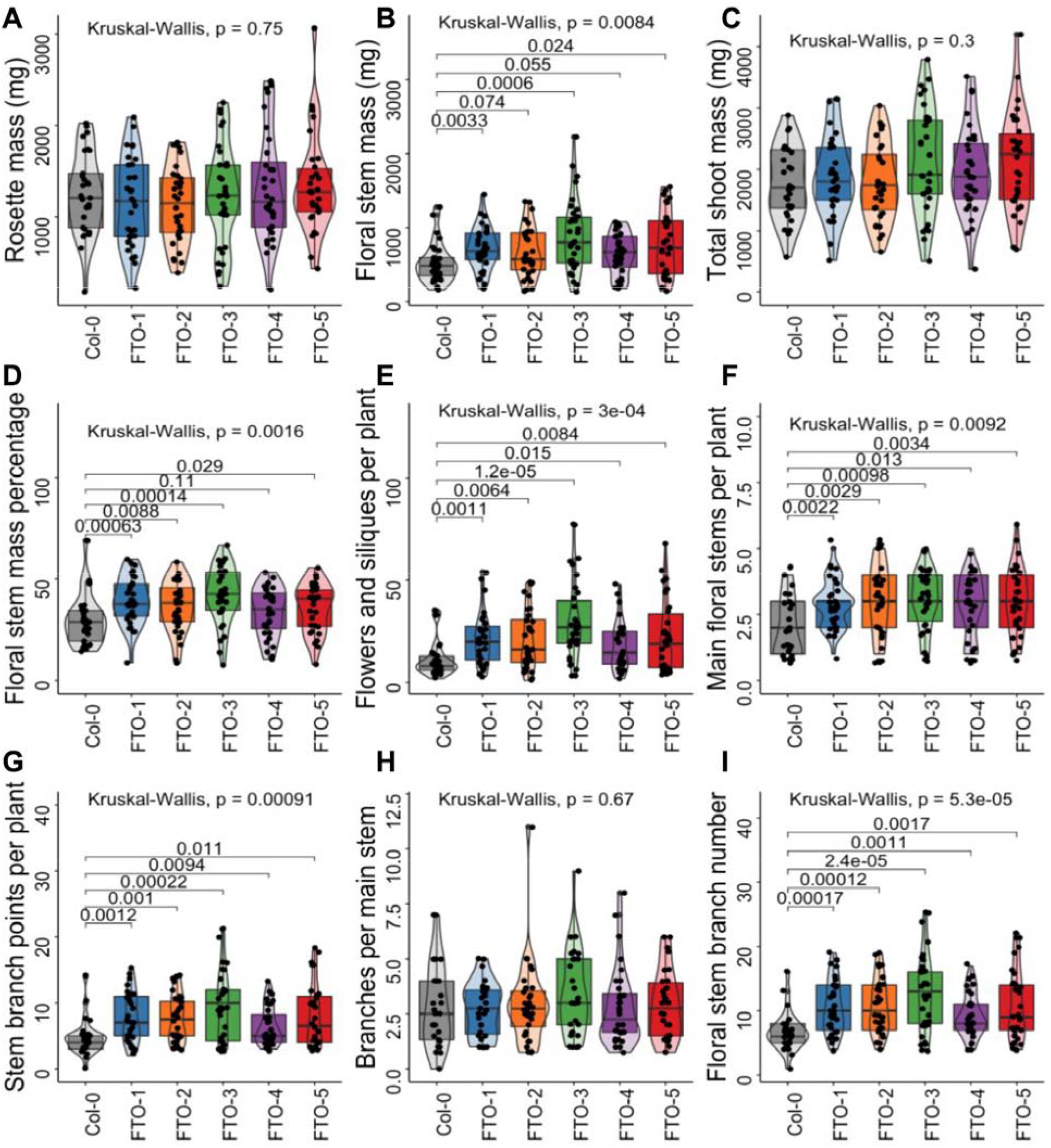
FTO increases many yield-associated traits in *Arabidopsis*. A) Rosette mass per plant. B) Floral stem mass per plant. C) Total shoot mass per plant. D) Floral stem mass percentage per plant. E) Number of flowers and siliques (fruits) per plant. F) Main floral stems per plant. G) Stem branch points per plant. H) Stem branch points per main stem. I) Floral stem branch number per plant. Boxplot central boxes cover the two central quartiles, points are raw data. Kruskal-Wallis p-value tests for significant difference between any of the groups, if Kruskal-Wallis result was significant then Wilcoxon test was performed between Col-0 and all FTO lines with Benjamini-Hochberg correction for multiple comparisons, pairwise p-values are displayed.

Since Yu *et al*. identify an approximately 40% increase in the tiller number as the primary driver of increased yield, we quantified the number of floral stems per plant and the number of floral branch stem points. The total number of floral stem branches increased by an average of 62% (Figure 2F). There was also a significant increase in the number of floral stem branches (Figure 2G), but the number of branches per main floral stem remained constant (Figure 2H). This indicates that the effect was driven by the increase in the number of main stems. The total number of floral stem shoot apical meristems – the sum of main stems and stem branch points – also showed a significant increase in all five lines (Figure 2I).

### FTO expression alters global gene expression patterns

We performed RNA-seq using RNA extracted from seedling shoot tissue to generate mechanistic hypotheses as to why transgenic expression of FTO might have these surprising effects. To ensure robustness of data with respect to experimental variability, we opted for six biological replicates, which enabled us to sequence Col-0 and three FTO independent lines, twenty-four samples in total. An average of 34 million reads were mapped per sample, gene-by-gene count data is available in Supplemental Table 2, raw sequencing data are available through the JGI portal (link in methods). In comparisons to Col-0, we found FTO-1 has far fewer differentially expressed genes than FTO-2 and FTO-3, both upregulated (Figure 2A) and downregulated (Figure 2B). Interestingly, FTO-1 also had a smaller phenotypic difference compared to Col-0 in floral stems per plant (Figure 2F). Given the smaller molecular and phenotypic effects observed for FTO-1, we decided to focus our analysis on the genes differentially upregulated in either all three lines or FTO-2 and FTO-3, leading us to 1436 upregulated and 1014 downregulated genes, loci and additional details of which are included in Supplemental Table 3.

We performed Gene Ontology (GO) term enrichment analysis using the PANTHER^26^ tool through TAIR, with the top 5 most enriched and least enriched (or most depleted) categories shown in Figure 3C and Figure 3D for the upregulated and downregulated gene lists, respectively. Additional GO categories along with their enrichment, number of genes, and significance can be found in Supplemental Table 4 and Supplemental Table 5 for upregulated and downregulated gene lists. For the upregulated gene list, the two most enriched categories are ‘protein maturation’ and ‘response to light intensity’, the two most depleted categories are ‘transmembrane transport’ and ‘defense response’. The next two upregulated categories both relate to porphyrin production, suggesting a change in chlorophyll metabolism. For the downregulated gene list, the most highly enriched GO categories were ‘proton motive force-driven mitochondrial ATP synthesis’ and ‘negative regulation of intracellular signal transduction’, and only the ‘unclassified’ category was depleted.

**Figure 3:**
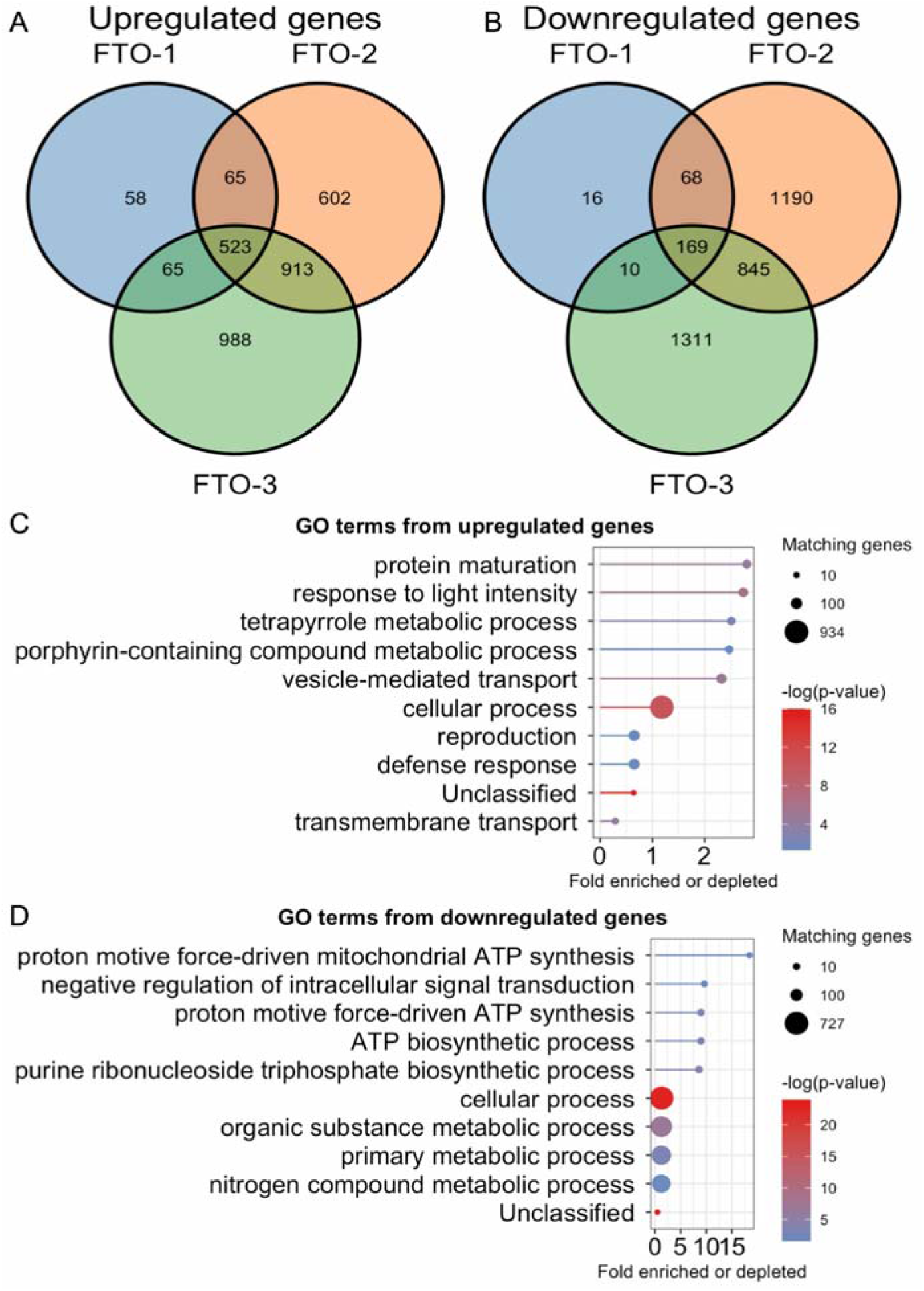
FTO expression alters global gene expression. A) Venn diagram of significantly upregulated genes in our three transgenic lines compared to Col-0. Multiple-comparison adjusted p-value threshold = 0.01. B) Venn diagram of significantly downregulated genes in our three transgenic lines compared to Col-0. Multiple-comparison adjusted p-value threshold = 0.01. C) Lollipop plot of Gene Ontology (GO) terms enriched among the upregulated genes. Dot size indicates number of genes, line length indicates fold enrichment, color indicates statistical significance. D) Lollipop plot of GO terms enriched among the downregulated genes. Dot size indicates number of genes, line length indicates fold enrichment, color indicates statistical significance.

### Growth-associated SAURs are overrepresented among the most highly upregulated genes

We identified the top 100 most upregulated and downregulated genes in each line to gain mechanistic insight into the function of FTO expression relative to Col-0. Similar to the broader search, we found there was more overlap between FTO-2 and FTO-3, and therefore chose to look at genes among the top 100 most-altered by fold change in either all three lines or just those two. This analysis yielded 41 shared most-upregulated genes and 36 shared most-downregulated genes (Supplemental Table 6). For both the highly upregulated and downregulated genes, there is much more overlap between lines than would be expected by chance (hypergeometric p-value = 2.53e-76 and 3.79e-64, respectively), confirming that the most highly-altered genes among all lines are strongly shared.

Among the 41 highly upregulated genes, 4 are SAUR genes which are known to regulate auxin-induced growth and development^27^. This is approximately 32-fold higher than expected based on frequency in the genome (hypergeometric p-value = 7e-06). Overexpression of SAUR genes increases plant growth^28,29^, and microRNA silencing reduces growth^29^. Specifically, we found SAUR10, SAUR28, SAUR63, and SAUR68 to be highly upregulated. SAUR10 has been shown to increase growth when overexpressed in *Arabidopsis*^30^, and was recently shown to be essential for normal growth and yield in rice^31^. SAUR28, along with three other nearby SAUR genes, was identified as the largest effect QTL for leaf architecture changes at bolting stage in response to different temperatures^32^. SAUR63 overexpression has also been shown to increase growth in *Arabidopsis* seedlings^33^. The increased expression of SAUR genes is particularly interesting in light of the reduced apical dominance and increased branching we observed and suggests that auxin regulation may be altered in FTO-expressing plants.

### Organellar genes and repressive transcription factors are overrepresented among the most highly downregulated genes

Among the 36 highly downregulated genes, we found 6 on the plastid and 6 on the mitochondrial genome, a ∼37-fold and ∼78-fold overrepresentation, respectively (hypergeometric p-values = 1.2e-08 and 1.3e-10). In contrast, 0 of the 41 highly upregulated genes were encoded in an organellar genome. Furthermore, among our 1436 significantly upregulated genes there were 0 mitochondrial and 0 plastid genes, whereas there were 21 mitochondrial and 34 plastid genes among the 1014 downregulated genes. This explains the enrichment of two different ‘proton motive force-driven ATP synthesis’ GO terms among our downregulated genes. Yu *et al*.’s claim that FTO expression increases nuclear genome-derived mRNA production offers one explanation, because one would expect to see a relative decrease in the abundance of mRNAs derived from organellar genomes. Yu *et al*. do not report RNA-seq data for organellar genes, and it should be noted that our library prep process using polyA selection should cause significant depletion of organellar compared to nuclear genome-derived transcripts, though there is no reason to expect this would differentially affect upregulated versus downregulated genes.

We also noticed several of the highly downregulated genes were transcription factors with known repression activity, including ERF13^34^, ERF22^35^, ZAT7^36^ and ZAT10^37^. Interestingly, *erf22* mutants have approximately 5-fold increased expression of SAUR10^38^, which concords well with our finding that FTO plants with highly reduced expression of ERF22 also have highly increased expression of SAUR10. Overexpression of ERF13^39^, ZAT7^36^ and ZAT10^40^ inhibit growth in *Arabidopsis*, and loss-of-function mutants of *erf22*^38^ and RNAi-silenced ERF13^41^ lines have enhanced growth rates. Taken together, these growth-reducing genes among the highly downregulated genes provide a plausible mechanism for the growth-enhancing effects of FTO. Downregulation of transcriptional repressors is also consistent with the claim by Yu *et al*. of an overall increase in nuclear-derived mRNA.

### Genes expressed in senescent tissue and genes associated with defense are overrepresented among the genes downregulated by FTO

As another method of exploratory data analysis, we used the Klepikova atlas^42^ through TAIR to identify the tissues in which each gene was most highly expressed. Of the 41 highly upregulated genes, 2 were most highly expressed in a senescent tissue (senescent leaf petiole and senescent leaf vein). In contrast, 11 of 36 highly downregulated genes were most highly expressed in a senescent tissue (all in senescent leaf petiole), a significant difference (Fisher’s exact test two-tailed p-value = 0.0046). Interestingly, all four repressive transcription factors were most highly expressed in senescent leaf petioles. It’s possible the downregulation of many genes expressed in senescent tissue is connected to the growth-enhancing phenotype.

We also collated GO terms for all genes, and noticed that 0 of 41 highly upregulated genes and 4 of 36 highly downregulated genes included ‘pathogen’ or ‘fungus’ in their GO terms, a significant difference (Fisher’s exact test two-tailed p-value = 0.044). Mutants of one of these genes, AT3-MMP, have been shown to be more vulnerable to the fungus *Botrytis cinerea*^43^. This supports our PANTHER GO-term analysis which found ‘defense response’ to be depleted among the upregulated genes (Figure 3C), and found the categories ‘defense response to bacterium’, ‘regulation of defense response’, and ‘defense response to fungus’ to all be enriched at least 2.3-fold among the downregulated genes (Supplemental Table 5). This reduction in expression level of pathogen-responsive genes is to be expected in light of long-established tradeoff between growth and defense^44^.

### Genes upregulated and downregulated by FTO expression differ in sequence properties

In general, RNA-seq data is best understood by considering genes and transcripts as units of biological function, which lends itself to methods of analysis like gene ontology and comparison of differentially expressed genes against other studies of the same gene. However, transcripts are also units of biological structure, and differences in structure such as length or nucleotide composition may be differentially affected by FTO expression. As such, we plotted the average nucleotide frequency within each transcript for three sets of genes: our 1436 upregulated genes, 1014 downregulated genes, and a set of 1200 genes chosen at random from our RNA-seq counts table as a reference point. The gene lists and sequences are available as Supplemental Table 3. We found the frequency of adenosine was significantly higher among our upregulated genes compared to both random and downregulated gene lists, with the majority of the gap being between upregulated and the other two categories (Figure 4A). Because m^6^A modification is primarily destabilizing, it stands to reason that with a transcriptome-wide reduction of m^6^A, we would expect to see adenosine-rich transcripts more often upregulated.

**Figure 4:**
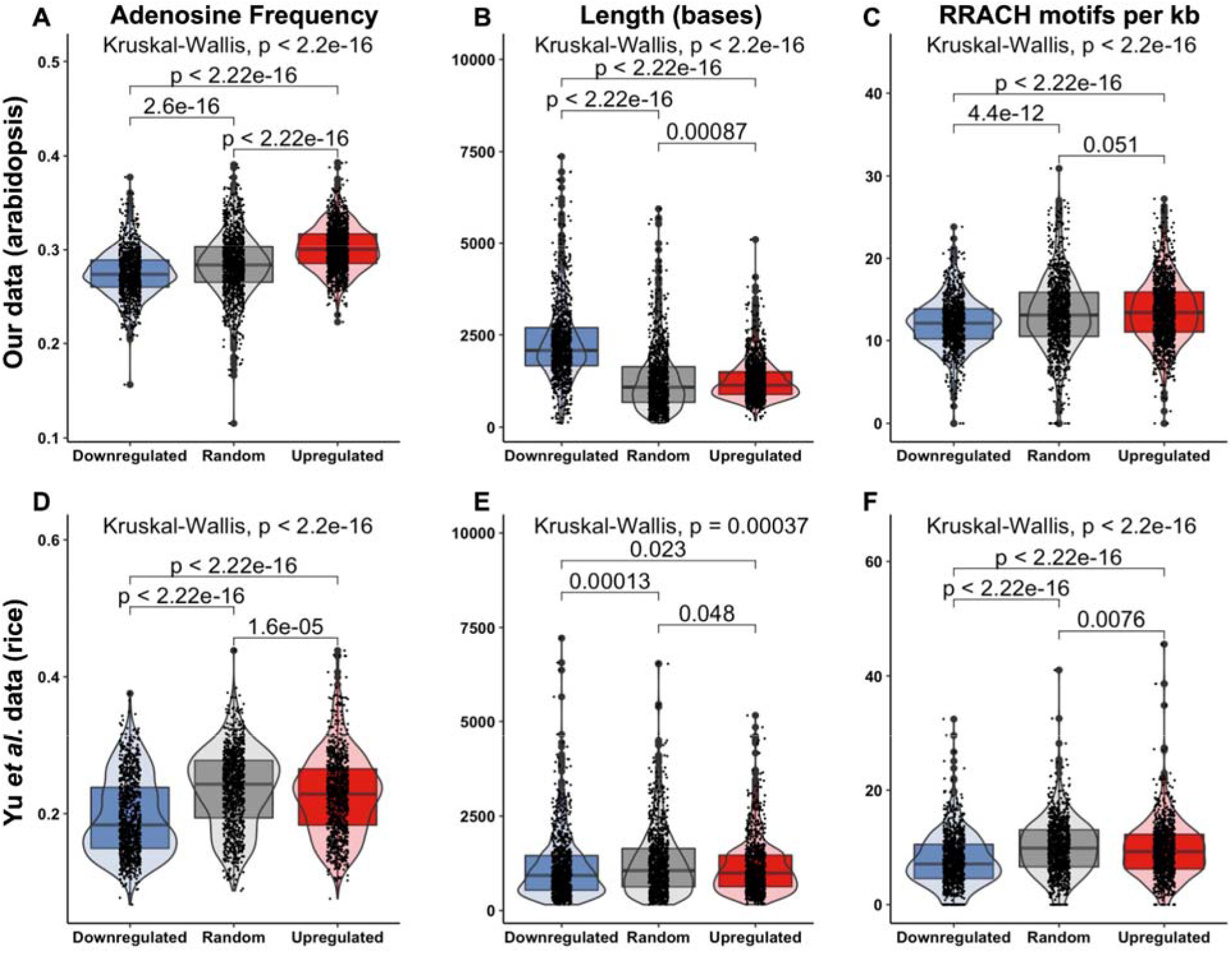
CDS composition of genes upregulated and downregulated by FTO expression are physically distinct. A) Frequency of adenosine in the CDS of transcripts downregulated and upregulated by FTO expression in our dataset. B) Length of the CDS of transcripts downregulated and upregulated by FTO expression in our dataset. C) Frequency of the RRACH (R = A/G, H = A/U/C) motif in the CDS of transcripts downregulated and upregulated by FTO expression in our dataset. D) Frequency of adenosine in the CDS of transcripts downregulated and upregulated by FTO expression in Yu *et al*.’s dataset. E) Length of the CDS of transcripts downregulated and upregulated by FTO expression in Yu *et al*.’s dataset. F) Frequency of the RRACH (R = A/G, H = A/U/C) motif in the CDS of transcripts downregulated and upregulated by FTO expression in Yu *et al*.’s dataset. Boxplot central boxes cover the two central quartiles, points are raw data. Kruskal-Wallis p-value tests for significant difference between any of the groups, if Kruskal-Wallis result was significant then Wilcoxon test was performed between all three categories of transcript with Benjamini-Hochberg correction for multiple comparisons, pairwise p-values displayed.

We next tested whether the length of transcripts differed in length between our three categories, and found the downregulated transcripts were much longer than either the random or upregulated transcripts: median length 90% and 81% longer, respectively (Figure 4B). Because m^6^A is known to localize to the motif RRACH (R = A/G, H = A/U/C), we next examined the frequency of that motif within each category of transcripts. Because transcript length differed, we normalized by length, and found that downregulated genes had fewer RRACH motifs per kb (Figure 4C). With the naïve assumptions of equal base frequency and random sequence, the RRACH motif occurs about 12 times per kb, among transcribed genes in *Arabidopsis* we found a slightly higher but comparable frequency. However, the majority of RRACH motifs are not methylated, in *Arabidopsis* the m^6^A frequency has been estimated at 0.5-0.7 per kb^3^. If there is a transcriptome-wide reduction in a destabilizing epitranscriptomic mark at RRACH motifs, we would expect relative enrichment of transcripts with a high concentration of those motifs among the upregulated genes. Genes whose transcripts are marked with m^6^A have been previously reported to be longer than genes whose transcripts are not methylated^45^. FTO-mediated removal of m^6^A and destabilization of those transcripts would cause an over-representation of longer genes among the downregulated gene list. We also tested the adenosine frequency, length, and RRACH motif concentration in the 5’ and 3’ UTRs and found the differences were either much smaller or nonexistent, it seems to be the CDS in particular that displays these striking structural differences between our different categories of transcript.

### Trends identified in our dataset are also found in previously-published RNA-seq datasets of plants with altered m^6^A deposition

We next investigated whether these structural differences between downregulated, randomly-selected, and up-regulated genes were present in previous RNA-seq datasets of m^6^A-altered plants. We begin with Yu *et al*.’s rice shoot RNA-seq dataset, which we used to generate even-sized groups of downregulated, randomly-selected, and upregulated genes in the FTO plants. We found genes upregulated by FTO to have higher CDS adenosine frequency (Figure 4D), matching our results and suggesting commonalities in the mechanism of action between rice and *Arabidopsis*. Median CDS length is only negligibly different between upregulated and downregulated genes in their dataset, though much like our dataset the upregulated gene list had fewer very long genes (Figure 4E), which is not a result of different size gene sets due to our balanced gene lists. Similar to our dataset, downregulated genes were depleted for RRACH motifs compared to randomly-selected or upregulated genes (Figure 4F). We performed GO analysis using PANTHER on the upregulated and downregulated gene lists, the results of which are available as Supplemental Table 7 and Supplemental Table 8, respectively. Genes associated with ‘response to light intensity’ are over-represented among their upregulated gene list. ‘Defense response’, ‘defense response to other organism’, and ‘response to fungus’ are all overrepresented among their downregulated gene lists, matching our results. For both gene lists, the only underrepresented GO term was ‘unclassified’.

To better understand how our results match previous *Arabidopsis* data, we also analyzed a publicly available RNA-seq dataset from *fiona1* mutant plants ^20^. We chose mutants of this specific m^6^A writer because the mutants of most others have severe phenotypes, whereas *fiona1* plants grow to maturity and comparable size to wild-type plants. We performed GO analysis using PANTHER after identifying upregulated and downregulated genes between Col-0 and *fiona1* plants, the results of which are available as Supplemental Table 9 and Supplemental Table 10, respectively. We found ‘defense response to bacterium’, ‘defense response to other organism’, and ‘defense response’ were all enriched at least 1.85-fold among the downregulated genes, whereas no defense categories were either depleted among the downregulated genes or enriched among the upregulated genes. The organellar genes were also non-evenly distributed between upregulated and downregulated genes in *fiona1*, among the upregulated genes there were 10 plastid and 1 mitochondrial gene, whereas among the downregulated genes were 40 mitochondrial and 1 plastid gene. The strong enrichment of mitochondrial genes in the downregulated gene list matches our findings with FTO, whereas the distribution of plastid genes is opposite to our finding.

We investigated several additional published RNA-seq datasets specifically for the question of organellar genes being overrepresented among the downregulated genes, and did not find a clear pattern — microRNA knockdown of the m^6^A writer MTA^18^ revealed in an overrepresentation of mitochondrial genes among the downregulated genes, but plastid genes were overrepresented among upregulated genes, matching the *fiona1* results. Mutants of the m^6^A writer FIP37^18^ show an enrichment for plastid genes among upregulated genes, whereas mitochondrial genes show no enrichment. Mutant plants deficient for the native m^6^A eraser ALKBH10B also show neither enrichment nor depletion for plastid or mitochondrial genes^22^.

## Discussion

Our findings broadly support the findings of Yu *et al*., demonstrating how the FTO overexpression approach may be a broadly transferable trait and should encourage future research using modulation of RNA methylation status as a tool to improve plant yield. We find a significant increase in flower and fruit number, stem branching, and reproductive stem biomass. We present several potential hypotheses for the effect, such as the upregulation of SAUR genes and repression of repressive transcription factors. We also present two striking observations for further investigation: an inconsistent but strong pattern of disproportional representation of organellar genes among the lists upregulated and downregulated by FTO, and differences between upregulated and downregulated gene sets such as a reduction in adenosine frequency and RRACH motif abundance among downregulated genes. We believe our data in conjunction with that of Yu *et al*. are sufficient to warrant more widespread research into the effects of FTO expression on plant yield and present directions for future research.

Public opinion has been a major barrier to the widespread deployment of transgenic crops, and we recognize that the introduction of a transgene derived from humans into crops represents a significant public relations liability. To hedge these concerns, future studies pursuing FTO overexpression may benefit from avoiding the human version of the gene, and instead utilize orthologs from close mammalian relatives such as porcine or bonobo orthologs. It may also be possible to achieve similar growth improvements with a transgene-free approach by reducing the level of RNA methylation through selective reduction of m^6^A writers rather than the addition of a phylogenetically-distant m^6^A eraser. However, the growth defects in reported m^6^A writer mutants suggest this will need to be approached carefully, and will likely require more mechanistic understanding of the basis of the growth enhancement phenotype. Fortunately, new sequencing technologies are rapidly improving our ability to detect RNA modification^46^, which may light the way towards knowing which RNA species in which tissues at which developmental stages are involved in this remarkable growth-enhancing phenotype.

## Supplemental files

Supplemental Table 1: Phenotypes of T3 generation plants

Supplemental Table 2: RNA-seq counts table

Supplemental Table 3: Gene IDs, sequence, and other parameters for upregulated, downregulated, and randomly-selected gene lists

Supplemental Table 4: PANTHER GO Term analysis of upregulated genes

Supplemental Table 5: PANTHER GO Term analysis of downregulated genes

Supplemental Table 6: Highly differentially expressed genes

Supplemental Table 7: PANTHER GO Term analysis of upregulated genes from Yu *et al*.’s dataset

Supplemental Table 8: PANTHER GO Term analysis of downregulated genes from Yu *et al*.’s dataset

Supplemental Table 9: PANTHER GO Term analysis of upregulated genes from publicly-available *fiona1* dataset

Supplemental Table 10: PANTHER GO Term analysis of downregulated genes from publicly-available *fiona1* dataset

## Methods

### Plant growth

Col-0 *Arabidopsis* were grown to flowering stage for transformation by floral dip and transformed with *Agrobacterium tumefaciens* strain GV3101 following standard methods, and the T0 progeny seeds selected for resistance to 50 mg/L kanamycin during axenic growth on agar plates. Over 50 independent lines were obtained, which were selected and propagated to increase seed and prepare for the main growth experiments with 5 lines of homozygous T3 plants. These plants were grown in a AR-95L3 chamber designed for *Arabidopsis* (Percival Scientific) under long day conditions with 12:12 light:dark, 25 °C, 60% relative humidity, “Professional Growing Mix” soil (Sungrow Horticulture) with Peter’s Professional 20-20-20 (ICL Growing Solutions) diluted into the water at 1 gram per Liter once per week. Plants were kept well watered in standard 1020 flats with 24 pots per flat.

### Drought experiments

We’ll see if this gets included

### Accession Numbers

Sequence data from this article can be found in the EMBL/GenBank database or the *Arabidopsis* Genome Initiative database under the following accession numbers: The human FTO enzyme used in this study is Genbank: KAI2578515, the porcine ortholog is GenBank: ADD69819. MTA is At4g10760, MTB is At4g09980, FIP37 is At3g54170, VIR is At3G05680, FIONA1 is AtMG00980, HAKAI is At5G01160, ECT2 is At3G13460, ECT3 is At5G61020, ECT4 is At1G55500, CPSF30 is At1G30460, and ALKBH10B is At4G02940. SAUR28 is At3G03830, SAUR10 is At2G18010. All other gene names are available in Supplemental Table 3. Raw sequencing data is available through the JGI portal: https://genome.jgi.doe.gov/portal/RNAofexinplants/RNAofexinplants.info.html

### RNA extraction and library preparation

15-30 mg of shoot tissue from day-old seedlings grown under axenic conditions on 1/2 MS plates was excised and immediately flash-frozen in liquid nitrogen and stored at -80 °C. RNA extraction was performed using E.Z.N.A Total RNA kit (Omega BioTek) following manufacturer directions. Extracted RNA was then DNAse treated with TURBO DNA-Free kit (Thermo Scientific). mRNA was isolated from an input of 1 ug of total RNA with oligo dT magnetic beads and fragmented to 300 bp - 400 bp with divalent cations at a high temperature. Using TruSeq stranded mRNA kit (Illumina), the fragmented mRNA was reverse transcribed to create the first strand of cDNA with random hexamers and SuperScript™ II Reverse Transcriptase (Thermo Fisher Scientific) followed by second strand synthesis. The double stranded cDNA fragments were treated with A-tailing, ligation with JGI’s unique dual indexed adapters (IDT) and enriched using 8 cycles of PCR. The prepared libraries were quantified using KAPA Biosystems’ next-generation sequencing library qPCR kit and run on a Roche LightCycler 480 real-time PCR instrument. Sequencing of the flowcell was performed on the Illumina NovaSeq sequencer using NovaSeq XP V1.5 reagent kits, S4 flowcell, following a 2x151 indexed run recipe.

### RNA-seq data analysis

#### Read preprocessing

Raw fastq file reads were filtered and trimmed using the JGI QC pipeline resulting in the filtered fastq file (*.filter-RNA.gz files). Using BBDuk [1], raw reads were evaluated for artifact sequence by kmer matching (kmer=25), allowing 1 mismatch and detected artifact was trimmed from the 3’ end of the reads. RNA spike-in reads, PhiX reads and reads containing any Ns were removed. Quality trimming was performed using the phred trimming method set at Q6. Finally, following trimming, reads under the length threshold were removed (minimum length 25 bases or 1/3 of the original read length - whichever is longer).

#### Read Alignment and Counting

Filtered reads from each library were aligned to the reference genome using HISAT2 version 2.2.1 (with -k 1 flag) [2] (BAMs/ directory). Strand-specific coverage bigWig files (fwd and rev) were generated using deepTools v3.1 [3] (bigWigs/ directory). featureCounts [4] was used to generate the raw gene counts (counts.txt) file using gff3 annotations. Only primary hits assigned to the reverse strand were included in the raw gene counts (-s 2 -p --primary options). Raw gene counts were used to evaluate the level of correlation between biological replicates using Pearson’s correlation and determine which replicates would be used in the DGE analysis (replicate_analysis.txt, replicate_analysis_heatmap.pdf). In the heatmap view, the libraries were ordered as groups of replicates. The cells containing the correlations between replicates have a purple (or white) border around them. FPKM and TPM normalized gene counts are also provided (fpkm_counts.txt and tpm_counts.txt).

#### Strandedness

Features assigned to the forward strand were also tabulated (-s 1 -p --primary options). Strandedness of each library was estimated by calculating the percentage of reverse-assigned fragments to the total assigned fragments (reverse plus forward hits).

#### Differential Gene Expression

DESeq2 (version 1.30.0) [5] was used to determine which genes were differentially expressed between pairs of conditions. The parameters used to call a gene differentially expressed between conditions were adjusted p-value < 0.01. The file DGE_summary.txt includes the log2 fold change, adjusted Pval and whether the gene is significantly differentially expressed (TRUE/FALSE/NA) for each pair of conditions specified. We also include shrunken log2 fold changes from DESeq2’s “normal” method for visualization and ranking. Refer to the Sample Summary Table above to match SampleName with the ConditionNumber assigned for analysis. Individual results for each pairwise comparison are in the directory Pairwise DGE Results. Note: Raw gene counts (counts.txt), not normalized counts are used for DGE analysis. DESeq2 conducts its own internal normalization using a sophisticated sampling model.

#### Gene Set Enrichment Analysis

Gene set enrichment analysis evaluates whether certain gene sets or pathways contain more differentially expressed genes than expected by chance. If KEGG annotations were available for the genes in this reference genome, PADOG (version 1.36.0) [6] was used to calculate gene set enrichment of the KEGG pathways found in the genome. Gene annotations are summarized in gene_functional_annotations.tsv and KEGG pathways with at least three genes present in the reference genome are provided in KEGG_gene_sets.tsv. Figures were produced using RStudio with the Tidyverse package ^47^.

#### Analysis of Yu *et al*. dataset

We used the shoot standard RNA-seq (as opposed to m^6^A-seq) data, and sorted genes by average fold-change between wild-type and FTO plants. We then generated our three gene lists by selecting the 1000 most-upregulated genes, 1000 most-downregulated genes, and 1000 genes selected using a random number generator.

## Acknowledgements

We acknowledge support from the DOE Joint BioEnergy Institute (http://www.jbei.org) supported by the U. S. Department of Energy, Office of Science, Office of Biological and Environmental Research, through contract DE-AC02-05CH11231 between Lawrence Berkeley National Laboratory and the U.S. Department of Energy. The work (proposal: 10.46936/10.25585/60008943) conducted by the U.S. Department of Energy Joint Genome Institute (https://ror.org/04xm1d337), a DOE Office of Science User Facility, is supported by the Office of Science of the U.S. Department of Energy operated under Contract No. DE-AC02-05CH11231. We thank Mitch Thompson and Simón Alamos for discussions during the preparation of this manuscript. We thank Daniel Peterson for comments and suggestions on our RNA-seq data analysis. We thank Peter Mellinger for watering plants.

## Author contributions

KM conceived and performed the experiments and wrote the manuscript. LW assisted with bioinformatic analysis. PS supervised and provided funding. All authors have read and approved the manuscript.

